# An intact S-layer is advantageous to *Clostridioides difficile* within the host

**DOI:** 10.1101/2022.11.22.517470

**Authors:** Michael J. Ormsby, Filipa Vaz, Joseph A. Kirk, Anna Barwinska-Sendra, Jennifer C. Hallam, Paola Lanzoni-Mangutchi, John Cole, Roy R. Chaudhuri, Paula S. Salgado, Robert P. Fagan, Gillian R Douce

## Abstract

*Clostridioides difficile* is responsible for substantial morbidity and mortality in antibiotically-treated, hospitalised, elderly patients, in which toxin production correlates with diarrhoeal disease. While the function of these toxins has been studied in detail, the contribution of other factors, including the paracrystalline surface layer (S-layer), to disease is less well known. Here, we highlight the essentiality of the S-layer *in vivo* by reporting the recovery of S-layer revertants, following infection with the S-layer-null strain, FM2.5. Sequencing of the *slp*A gene revealed either correction of the original point mutation or modification of the sequence upstream of the mutation, which restored the reading frame, and translation of *slpA*. Selection of these strains was rapid, with up to 90% of isolates identified as revertants 24 h post infection.

Two revertant isolates, RvA and RvB, showed modification of 3 and 13 amino acids respectively, compared to wild type sequence. Structural determination of SlpA from RvB revealed a different orientation of its domains, resulting in a reorganisation of the lattice assembly and changes in interacting interfaces which might result in functional differences. These revertants showed differing patterns of disease i*n vivo*; RvA causing equivalent severity to R20291 and RvB an attenuated FM2.5-like phenotype. Comparative RNA sequencing (RNA-Seq) analysis of *in vitro* grown isolates showed large changes in differentially expressed genes (DEGs) between R20291 and FM2.5 namely in TcdA/TcdB expression, in transcripts associated with sporulation and those linked to cell wall integrity, which may account for attenuation observed *in vivo*. In comparison, smaller differences were observed between RvA/R20291, and RvB/FM2.5 respectively, which correlated with observed disease severity *in vivo*. Cumulatively, these data highlight that the S-layer plays a role in *C. difficile* disease.

**Author Summary:** The S-layer of *C. difficile* is a paracrystalline array that covers the outer surface of the bacterial cell but its contribution to overall disease remains unclear. A previously described, spontaneous *slpA*-null mutant, FM2.5, with a point mutation in *slp*A offered an opportunity to study the role of the S-layer *in vivo*. Here, we confirm that this strain is less virulent *in vivo* despite effectively colonising the host and producing toxin. We also show *in vivo* selection for sequence modifications that restore *slp*A translation and produce an S-layer. While such modifications do not affect the overall 3D structure of individual SlpA (sub)domains, they can lead to altered orientation of the structural domains and subsequent S-layer assembly. Importantly, RNA-Seq analysis *in vitro* showed large differences in gene expression between FM2.5 and R20291. Detected differences in transcription of genes involved in toxin expression and sporulation suggests that the S-layer provides a selective survival advantage within the host, which contributes to disease severity.

## Introduction

*Clostridioides difficile* is the most common cause of hospital acquired diarrhoea globally, with disease linked to disruption of the intestinal microbiota through antibiotic use (Smits et al. 2016). The virulence of *C. difficile* has widely been attributed to the production of two toxins; toxins A (TcdA, enterotoxin) and B (TcdB, cytotoxin), responsible for cytoskeletal modifications, epithelial damage, inflammation, and fluid loss (Braun et al. 1996; Chandrasekaran and Lacy 2017). A third toxin, the binary *C. difficile* toxin (CDT), expressed by only a subset of strains, has been linked to enhanced disease severity (Chandrasekaran and Lacy 2017). Consequently, *C. difficile* colitis has widely been considered as a toxin-mediated disease. However, the availability of tools to analyse gene expression and improved methods of mutagenesis (Cartman et al. 2012a), together with the availability of an accessible murine animal model (Chen et al. 2008a; Theriot et al. 2011), have offered new opportunities to identify other traits, both bacterial and host-associated, that impact disease severity (McDermott et al. 2017; Fletcher et al. 2021). The recent use of such approaches has provided clearer understanding of the metabolic flexibility of these organisms, the role of the microbiome in disease progression (Buffie et al. 2015; Fletcher et al. 2021; Girinathan et al. 2021) and established several bacterial factors that influence the host response (Maldarelli et al. 2014; Batah et al. 2017; Arato et al. 2019). Of particular interest in this context, is the role of the S-layer in disease. This paracrystalline protein array is the outermost layer of the *C. difficile* cell envelope, with similar structures found in many bacteria and virtually all archaea (Fagan and Fairweather 2014).

The S-layer has been shown to perform multiple and vital roles including providing protection from environmental factors such as variations in pH, mechanical and osmotic stresses (Engelhardt and Peters 1998; Claus et al. 2002; Engelhardt 2007). *In vivo*, it is proposed to play a role in molecular sieving (Sleytr and Beveridge 1999) and ion trapping, protecting the organism from antimicrobial peptides and bacteriolytic enzymes produced in response to infection (Lortal et al. 1992; Kirk et al. 2017). The S-layer has also been shown to be a key target in bacteriophage predation (Callegari et al. 1998; Kirk et al. 2017; Royer et al. 2022).

In *C. difficile*, the main component of the S-layer is SlpA, which is post-translationally cleaved by a cell wall protein (CWP), Cwp84, into two functional S-layer proteins (SLPs), SLP_L_ and SLP_H_ (Kirby et al. 2009; Fagan and Fairweather 2014). The proteinaceous array is further decorated by other CWPs, which provide additional functionality (Fagan and Fairweather 2014). Assembly of the paracrystalline array relies on tiling of SLP_H_ triangular prisms on the cell wall, interlocked by SLP_L_ ridges facing the environment (Lanzoni-Mangutchi et al. 2022). Exposure of SLP_L_ to the environment is consistent with its high sequence variability observed between different *C. difficile* strains, with 13 different S-layer cassette types (SLCTs) identified to date (Dingle et al. 2013; Kirk et al. 2017). Strikingly, the lattice is very compact compared to other studied S-layers, which have pores of between 30 – 100 Å compared with only ^~^10 Å in diameter in *C. difficile* (Lanzoni-Mangutchi et al. 2022). This tight packing correlates well with the hypothesis that S-layer acts as a molecular sieve (Sleytr and Beveridge 1999), as deletion of the most exposed regions of SLP_L_ results in a strain with increased sensitive to lysozyme, in comparison to the parent strain, R20291 (Lanzoni-Mangutchi et al. 2022).

In *C. difficile*, the S-layer has also been implicated in host cell adhesion (Merrigan et al., 2013), biofilm formation (Kirby et al. 2009; Dapa et al. 2013; Richards et al. 2018) and immunomodulation through cell signalling of the host response (Ausiello et al. 2006; Sakakibara et al. 2007; Sekot et al. 2011). SlpA has been shown to induce innate and adaptive immune responses through activation of TLR4 (Ryan et al., 2011). However, the role of the S-layer in *C. difficile* pathogenesis and in immune evasion remains poorly understood.

Previously, we reported the isolation and characterization of a spontaneous *C. difficile* strain lacking an S-layer, FM2.5 (Kirk et al. 2017). In initial studies using the Golden Syrian hamster as the infection model, FM2.5 caused no symptoms of disease, despite effectively colonising infected animals (Kirk et al. 2017). However, the acute sensitivity of hamsters to *C. difficile* toxins and lack of readily available immunological tools, limits their usefulness in studying the more nuanced facets of this infection. In contrast, mice are naturally less susceptible to CDI, requiring more extensive antibiotic treatment to suppress the flora, and higher challenge doses to achieve colonisation (Chen et al. 2008b; Winston et al. 2016). However, mice offer greater opportunities to determine the contributions of other virulence-associated traits on disease outcome, including long-term persistence associated with relapsing disease (Best et al. 2012).

Here, we sought to elucidate the role of the S-layer as a major virulence determinant in a murine model of infection, to determine whether the loss of virulence observed in the hamster model is reciprocal in other hosts. Our results suggest that the S-layer offers a competitive colonisation advantage within the mouse intestine and is important for *in vivo* disease severity.

## Results

### The S-layer contributes to severe disease in the murine model of *C. difficile*

In a murine model of infection, loss of body weight offers a strong correlative measure of *C. difficile* disease severity (Jukes et al., 2020). Infection of antibiotic pre-treated mice with strain R20291 resulted in significant weight loss of up to 15%; peaking between 24 and 48 h post-infection (hpi). In contrast, mice infected with the S-layer deficient derivative strain FM2.5 showed consistently less weight loss (average 6%; Fig. 1a). Measurement of total *C. difficile* in faecal material showed comparable levels of shedding at 24 and 72 hpi (Fig. 1b), while analysis of total *C. difficile* present in caecal (Fig. 1c) and colonic (Fig. 1d) material taken at post-mortem indicated a trend for less recovery of FM2.5 24 hpi. However, comparable levels of FM2.5 and R20291 were recovered from these tissues at 48 and 96 hpi. Analysis of *C. difficile* spores within the total faecal, caecal and colonic material showed a similar trend, with less FM2.5 spores recovered at 24 hpi, and comparable numbers of spores at 72 and 96 hpi (Fig. S1).

**Fig 1.**
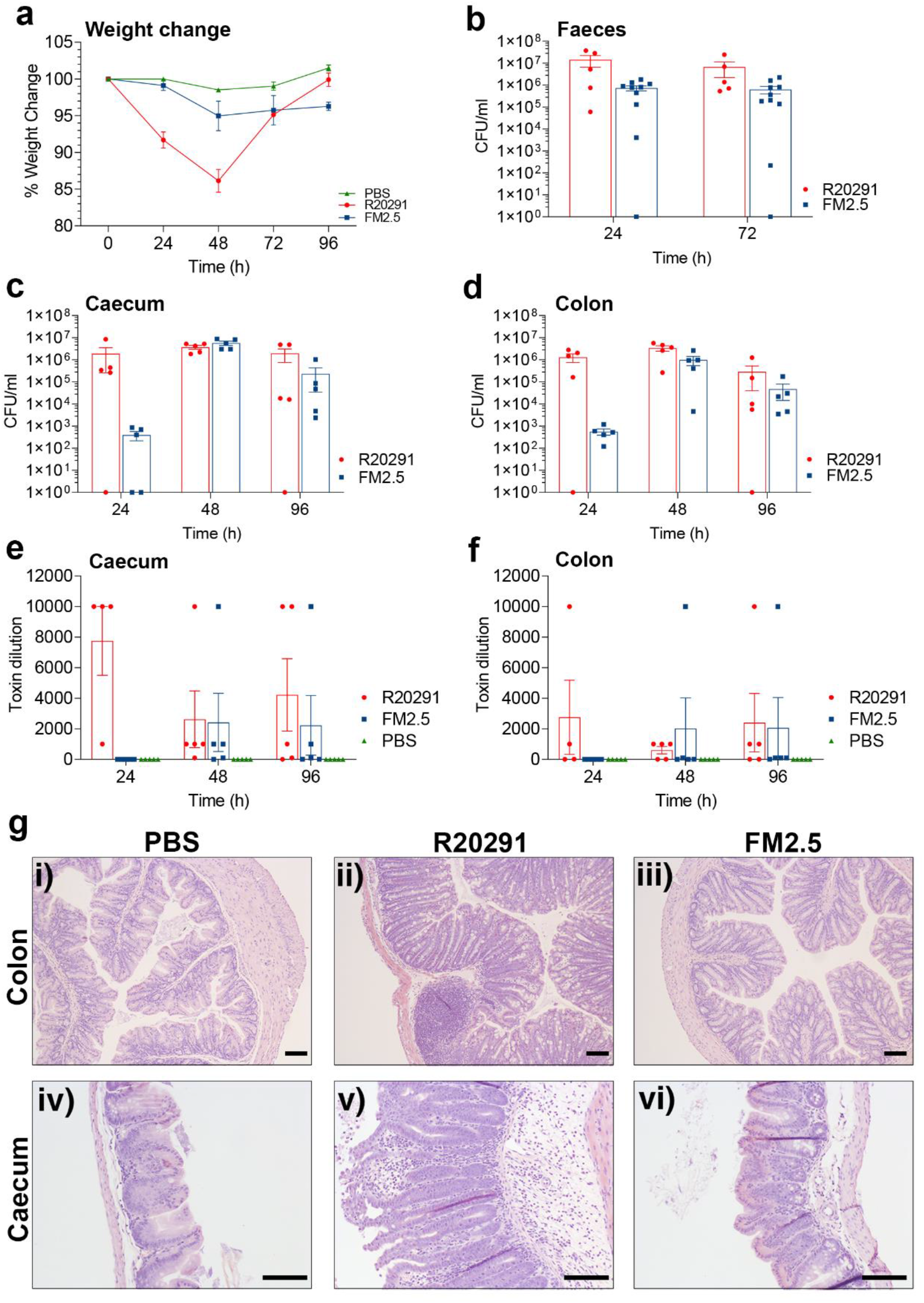
SlpA deficient *C. difficile* is less pathogenic in a murine model of infection. Female C57/Bl6 mice were challenged with spores of R20291 or FM2.5, or mock infected with sterile PBS. (**a**) Weight loss was monitored every 24h for four consecutive days following infection. Each point is the average of several replicate experiments (n>3), with at least 5 animals per time point. (**b**) CFU/ml of faecal material collected at 24 and 72 hpi. (**c**) CFU/ml of caecal content at 24, 48 and 96 hpi. (**d**) CFU/ml of colon content at 24, 48 and 96 hpi. (**e**) Toxin activity of caecal content and (**f**) colonic content was determined at 24, 48 and 96 hpi; through challenge of Vero cells *in vitro*. Results displayed indicate the reciprocal of lowest dilution at which toxin activity could be measured. (**g**) Histopathological sections representing colon (i, ii and iii) and caecal (iv, v and vi) sections following challenge with PBS (i and iv); R20291 (ii and v); or FM2.5 (iii and vi). Scale bars represent 100 μm. Results displayed are the mean ± SEM of at least three independent biological replicates. Statistical tests were conducted using GraphPad Prism software v.12. Statistical tests include one-way ANOVA with Tukeys post-test; or a student’s t-test with Welch’s correction. Statistical significance is indicated: ns – not significant; *p < 0.05; **p < 0.01; and ***p < 0.001.

Interestingly, mice infected with R20291 that survived infection showed full signs of recovery by 96 hpi, returning to pre-infection weights, equivalent to those of non-infected mice. In contrast, FM2.5-infected mice failed to return to their pre-infection weight even when animals were monitored for a further five days (9 days pi; typical profile Fig. 4c) despite the animals remaining asymptomatic, with no evidence of loose faeces.

Assessment of *in vivo* toxin production by both R20291 and FM2.5 in the caecum (Fig. 1e) and colon (Fig. 1f) revealed that, at 24 hpi, less toxin was recovered from mice infected with FM2.5. However, at 48 and 96 hpi, comparable levels of toxin were recovered. Histological examination (at 48 hpi) showed that mice infected with R20291 displayed significantly greater pathology than those infected with FM2.5 and a PBS challenged control group (Fig. 1g i-vi. Cumulative scoring of tissue damage is reported in Fig. S1).

### *In vivo* pressure drives selection for S-layer revertants

When grown on selective chromogenic agar (ChromID®; BioMerieux), the morphology of R20291 presents with the typical ‘fried egg’ *C. difficile* colony, visible after approximately 16 h of incubation (Fig. 2a). In contrast, FM2.5 produces smaller and smoother colonies, which take ^~^24 h to emerge (Fig. 2b). Following infection of mice with R20291 and FM2.5, faecal material was recovered and plated daily. During examination of resultant colonies, it was noted that, while colonies from R20291 infected mice showed the expected morphology, material retrieved from FM2.5-infected mice showed a mixture of both large (FM2.5_large_) and small colony types (FM2.5_small_) (Fig. 2b). Additionally, FM2.5_large_, were countable after 16 h incubation, while the expected FM2.5-like colonies, FM2.5_small_, were only observable from 24 h. Several colonies from both types, FM2.5_small_ and FM2.5_large_, were streaked from the original plates and were sub-cultured twice to ensure clonality. Individual clones were then stored at −80°C.

**Fig. 2.**
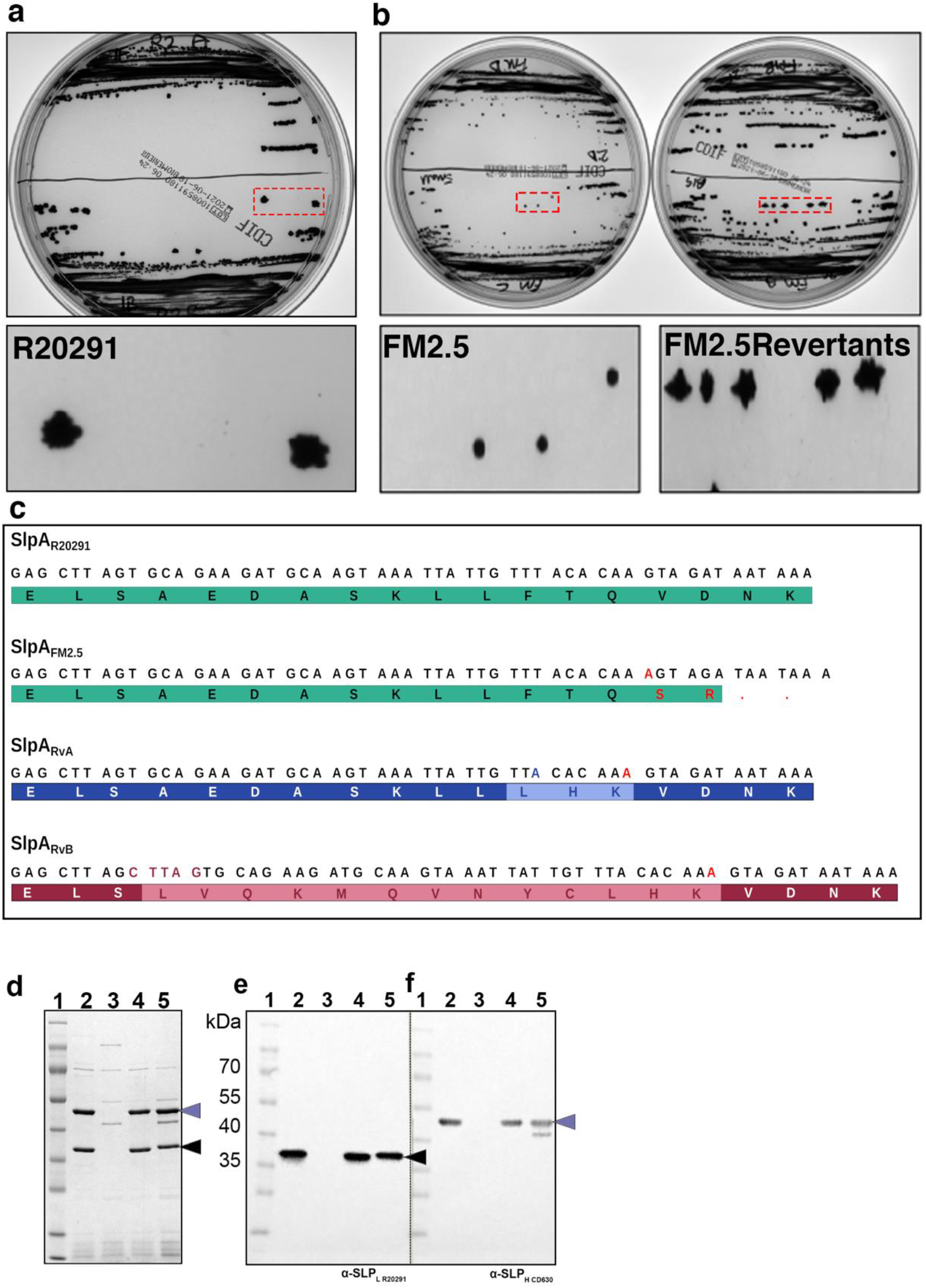
Recovery of S-layer revertants following *in vivo* challenge. Following challenge with spores of R20291 and FM2.5, faecal material was recovered and plated on *C. difficile* selective chromogenic agar (Biomerieux). (**a**) Colonies of R20291. The dashed red boxed area is enlarged and shown below. (**b**) The two colony types of FM2.5. The dashed red boxed area is enlarged and shown below. FM2.5_large_ is representative of the large colony phenotype recovered, while FM2.5_small_ is indicative of the typical FM2.5 colony morphology. (**c**) Sequencing of a region of *slpA* shows the sequence of R20291; the insertion (red) in FM2.5 responsible for the truncation of the SlpA protein; a single nucleotide deletion (blue) in the FM2.5 sequence, resulting in RvA; a five-nucleotide insertion (crimson) in FM2.5 sequence resulting in RvB. (**d**) SDS-PAGE analysis of cell wall proteins extracted by low pH preparation. Lane 1: MW Marker; Lane 2: R20291; Lane 3: FM2.5; Lane 4: RvA; Lane 5: RvB. (**e**) Western immunoblot analysis using an anti-SLP_H_ antibody. (**f**) Western immunoblot analysis using an anti-SLP_L_ antibody.

Amplification of the *slpA* sequence from these clones revealed that the FM2.5_large_ colonies contained modifications in the genomic sequence upstream of the FM2.5 mutation site (single nucleotide insertion, Fig. 2c). Several sequence variants were identified, with two most common mutants named Revertant A (RvA) and Revertant B (RvB). In RvA, a single nucleotide deletion (246delT) restored the original reading frame, rescuing translation of the full SlpA; modifying three amino acid residues in the translated protein. In, RvB, an insertion of five-nucleotides (249_253insCTTAG), which again restored the reading frame and resulted in modification of 13 amino acids within the mature protein (Fig 2c Interestingly, revertants were identified in several *in vivo* experiments, using batches of independently prepared spores.

To confirm expression of SlpA, low pH cell surface extracts of strains RvA and RvB were analysed. SDS-PAGE showed that both SLP_H_ and SLP_L_ proteins were present in R20291, absent in FM2.5 but restored in RvA and RvB (Fig. 2d). This was confirmed through western immunoblot analysis using anti-SLP_H_ and anti-SLP_L_ antibodies (Fig. 2e and f).

### Reversion can affect SlpA structure and assembly

To understand the effects of the detected reversions on SlpA structure and S-layer assembly, crystallisation of SlpA_R20291_, SlpA_RvA_ and SlpA_RvB_ was carried out. Although crystals were obtained for all three variants, only SlpA_RvB_ crystals were of sufficient quality for x-ray diffraction data collection and structural determination by molecular replacement, using previous SlpA structures as models, including a variant of SlpA_R20291_ lacking the most exposed region of SLP_L_ - SlpA_RΔD2_ - (PDB ID: 7ACZ) (Lanzoni-Mangutchi et al. 2022). SLP_H_ and the interacting domains were easily traceable in the electron density but D1 was only partially built, whilst density for domain D2 was very poor and this region could not be traced in the final SlpA_RvB_ model (Fig. 3a, PDB ID: 8BBY, Table S1). This implies that D2 is flexible and/or unstructured, while the structure of the core domains required for S-layer assembly - SLP_H_, and, to a lesser extent, D1 and LID/HID (Lanzoni-Mangutchi et al. 2022) - seems to be generally maintained. However, the relative orientation of these domains in the SlpA molecule is altered (Fig. 3a), with D1 and the interacting domains rotated towards the SLP_H_ plane by ^~^30 ° (Fig. 3a). The 13 altered residues in α2_L_ in SlpA_RvB_ result in disruption of the α-helix secondary structure and introduce disorder in the upstream loop that links the preceding β-strand (β3_L_) and α2_L_. It is worth noting that SLP_H_ in R20291, which belongs to SLCT 4, has several insertions within the cell wall binding 2 (CWB2) sequence motifs that define CWPs in *C. difficile*, when compared to other SlpA types. These insertions could not be traced in our SlpA_RΔD2_ model (Lanzoni-Mangutchi et al. 2022) but were traceable in the SlpA_RvB_ and result in several loops protruding above the SLP_H_ plane, towards the environment, partially occluding the CWB2 motifs (Fig. 3a, right). Together with the movement of the interacting domains and D1 towards the SLP_H_ tiles, this creates a more compressed arrangement (^~^66 Å compared to ^~^76 Å in SlpA_CD630_, PDB ID: 7ACY, Fig. 3b, bottom).

**Fig. 3.**
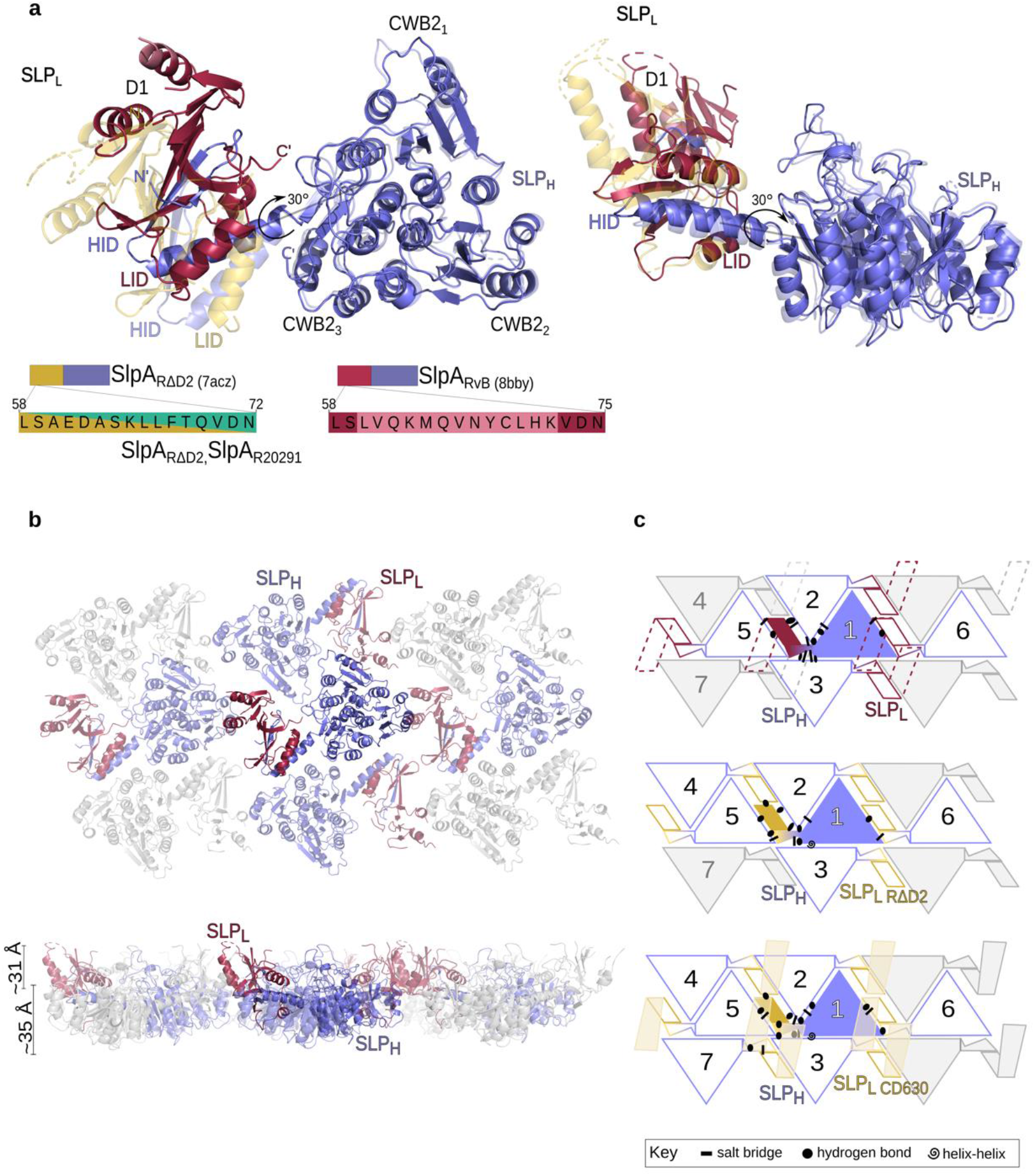
Structure of SlpA_RvB_ shows a different assembly arrangement. (**a**) Structural model of SlpA_RvB_ (SLP_L_ – pale red, SLP_H_ – slate blue, PDB ID: 8BBY), superimposed on SlpA_RΔD2_ model (SLP_L_ – gold, SLP_H_ – slate blue, semi-transparent), with the rotation angle of the D1 and LID/HID domains shown by an arrow. Three distinct structural features are observed: SLP_H_, LID/HID and D1. Cartoon representation of the SLP_H_/SLP_L_ (H/L) complex, as seen from the environmental side (left) and side view (right). Sequence of α2_L_, with paler colours indicating differences, is shown schematically. (**b**) Cartoon representation of the H/L planar array (PDB ID 8BBY, interacting molecules coloured and viewed as in **a**). (**c**) 2D schematic of H/L complex crystal packing in SlpA_RvB_ (top), SlpA_RΔD2_ (centre) and SlpA_CD630_ (bottom), indicating the interaction network linking a single H/L (slate blue/crimson or slate/blue/gold) complex with neighbouring molecules in a planar arrangement generated by SLP_H_ tiling. The missing D2 in the SlpA_RvB_ model is represented as dashed lines. Notably, D1-D1 interactions seen in other models are missing in RvB and the SLP_H_ tiles are shifted, with new HID-CWB2_3_ interactions stabilising the lattice. Array is depicted as seen from the extracellular environment, with symbols representing key interaction types in the crystal lattice, detailed in Table S2.

In the crystallographic models of SlpA_CD630_ (PDB ID: 7ACY), SlpA_R7404_ (PDB ID: 7ACX) and SlpA_RΔD2_ (PDB ID: 7ACZ), α2_L_ was responsible for closing a gap between neighbouring SlpA molecules via D1-D1 interactions (Lanzoni-Mangutchi et al. 2022). In the crystallographic model of the R20291-derived SlpA_RΔD2_ variant, D1-D1 interactions are mediated by hydrogen bonds between S50_L_-S50_L_ and Q70_L_-A49_L_ from neighbouring molecules.

In the SlpA_RvB_ structure, disruption of α2_L_ and reorientation of D1 and LID/HID relative to SLP_H_ leads to changes in the interactions between neighbouring molecules and, consequently, a rearrangement of the S-layer array. Strikingly, the D1-D1 interactions seen in previous models were not observed here, possibly due to the flexibility of α2_L_ and preceding loop caused by the changes in the reverted sequence leading to a different orientation of D1 domains. Unlike in the previously determined structures, neighbouring D1 domains in the SlpA_RvB_ structure are too far apart to mediate contacts (> 12 Å). In the previous models, SLP_H_ tiling creates two wide channels, which are stabilised by interactions mediated by the interacting domains and D1 (Lanzoni-Manguthi, 2022). A different mode of stabilising the SLP_H_ tiling is observed in SlpA_RvB_, with the interacting domains now partially inserted in those cavities (Fig. 3b, Fig. S2a). A new interacting interface between HID from one molecule and CWB23 occludes these gaps and stabilises the S-layer lattice (Fig. 3c, Fig. S2a). This new arrangement of the crystal lattice is in line with our proposed assembly model, where the S-layer 2D array is maintained mostly by hydrogen bonds and salt bridges across surfaces with complementary charges (Lanzoni-Mangutchi et al., 2022), largely dependent on SLP_H_-SLP_H_ interactions and stabilised by varying degree of interactions involving SLP_L_ (Fig. 3c and Table S2). The structural model of RvB confirms that changes in SLP_L_ can be accommodated with minor structural changes to the (sub)domains, by exploring flexible loops and hinges to provide a stable S-layer.

As no crystal data was obtainable, we also calculated models for SlpA_R2021_ and SlpA_RvA_ using the SWISS-MODEL server, based on previous models (Lanzoni-Mangutchi et al. 2022) and the SlpA_RvB_ structure determined here. Depending on which template was used (SlpA_RΔD2_ or SlpA_RvB_), different predicted structures of SlpA_RvA_ were obtained, varying mostly in the orientation of D1 and interacting domains relative to SLP_H_ (Fig. S2b). Interestingly, one common feature was that the changes resulting from the revertant sequence seem to be accommodated not by altering the α-helix but by varying the length of the upstream loop that links the preceding β-strand (β3_L_) and α2_L_ (Fig. S2b). It is therefore unclear if SlpA_RvA_ is more likely to adopt a R20291-like as observed in the SlpA_RΔD2_ model or RvB-like S-layer assembly, as both can accommodate the modified sequence.

### *In vivo* S-layer selection is independent of toxin expression

FM2.5 has previously been observed to show a delay in toxin production (Kirk et al. 2017), consequently we chose to investigate whether *slpA* reversion was accompanied by a potential restoration of toxin production *in vivo*. Mice were infected with FM2.5*ΔPaLoc*, in which the **<u>Pa</u>**thogenicity **<u>Loc</u>**us (PaLoc), encoding toxins A and B, had been deleted. In contrast to mice infected with FM2.5, animals challenged with FM2.5*ΔPaLoc* showed no weight loss over the 96 hours of infection (Fig. 4a) and as expected, no toxin was observed in the caecum or colon (Fig. S3c). Total bacteria (Fig. S3a) and spores (Fig. S3b) recovered from the caecum and colon of these mice were comparable at 96 hpi to that observed in animals infected with FM2.5.

**Fig 4.**
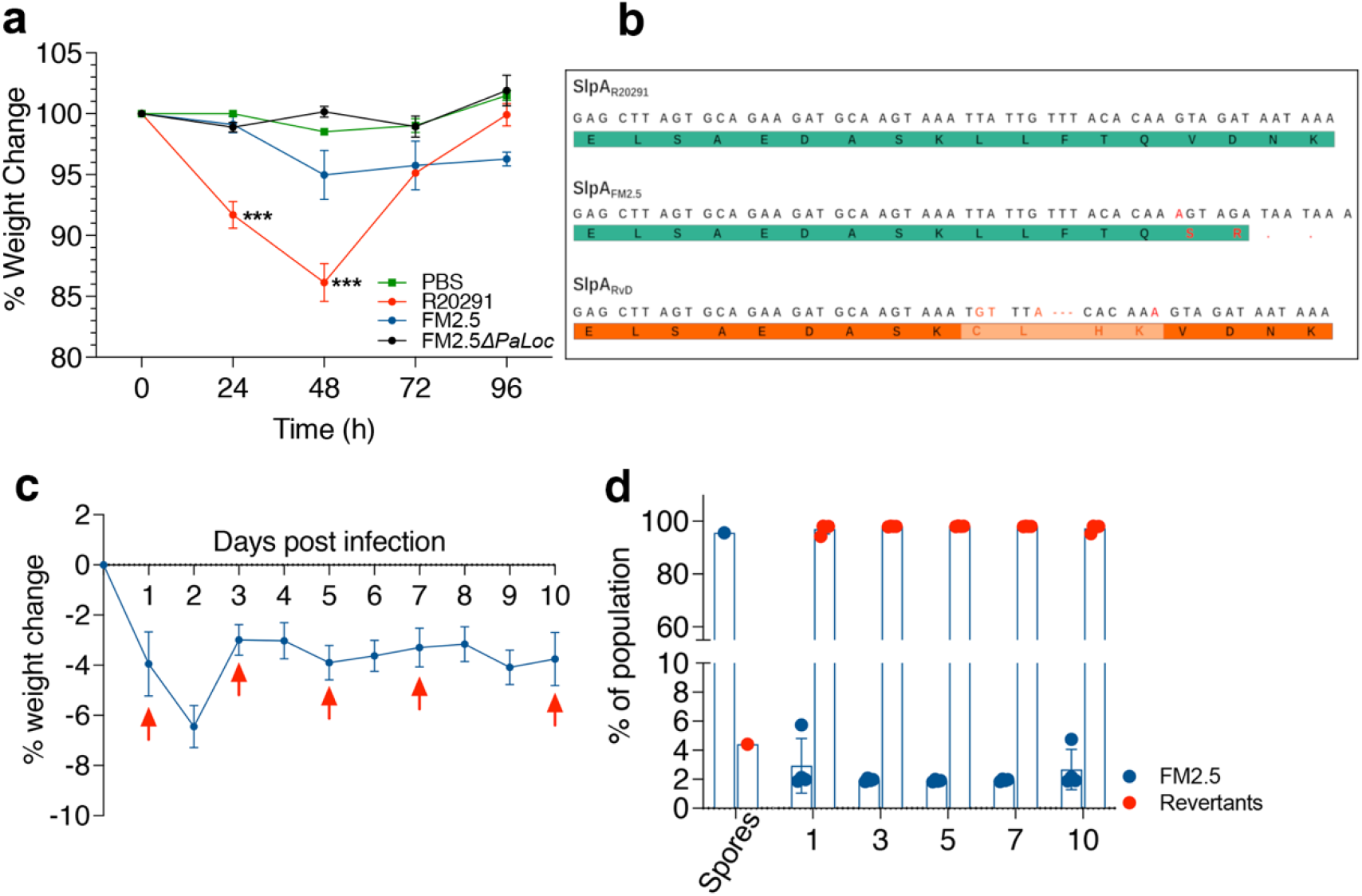
*In vivo* challenge of mice with FM2.5*ΔPaLoc*. Female C57/Bl6 mice were challenged with spores of R20291, FM2.5 and FM2.5*ΔPaLoc*. (**a**) Weight loss was monitored every 24h for four consecutive days following infection. Each point is the average of multiple mice, where n≥ 5. (**b**) Sequencing of a region of *slpA* shows the sequence of R20291; the insertion in FM2.5 (red) responsible for the truncation of the SlpA protein; with an additional revertant named RvD, which shows a complex array of sequence insertions and deletions (orange) in this region of the *slpA*. (**c**) Time course showing infection and weight loss in nine mice infected with FM2.5. Arrows indicate times of faecal sample collection used in amplicon sequencing of the variable region of *slpA*. (**d**) Description of relative proportion of FM2.5 sequences in samples analyzed prior to and post infection.

Interestingly, S-layer revertants were also recovered from these mice, as identified by sequence modifications in the same region of *slpA* compared to the R20291 sequence (RvD, Fig. 4a). These changes also facilitated restoration of an intact S-layer, indicating that any potential selection advantage is independent of toxin production.

Reproducible recovery of S-layer variants *in vivo* raised the possibility that low numbers of genetic variants exist within the FM2.5 population, which are amplified by the *in vivo* environment. To test this hypothesis, we undertook amplicon sequencing of *slpA* in the spore preparations used for mouse infections, and in bacteria recovered from faecal material from mice infected with FM2.5 at 24, 48, 72 and 96 hpi (Fig. 4c). This analysis revealed that revertants in which the original frameshift mutation found in FM2.5 was corrected by deletion of the extra nucleotide (252delA, RvC) were present in the initial spore preparation, albeit as a low proportion of the population (<5%, Fig. 4d). Isolation of revertants as early as 24h (>94% of the population) suggests that expressing an intact S-layer provides a competitive advantage *in vivo* over the S-layer deficient strain.

### The revertant strains display differing levels of virulence in mice

To assess whether recovery of SlpA by revertants correlated with rescued virulence, mice were infected with spore preparations of RvA and RvB, alongside R20291 and FM2.5 (Fig. 5a). Interestingly, infection with RvA resulted in significant weight loss within the first 48h of infection, which was similar to mice infected with R20291. RvB, in contrast, showed a similar limited pattern of weight loss to animals infected with FM2.5, which stabilized from 48 hpi.

**Fig 5.**
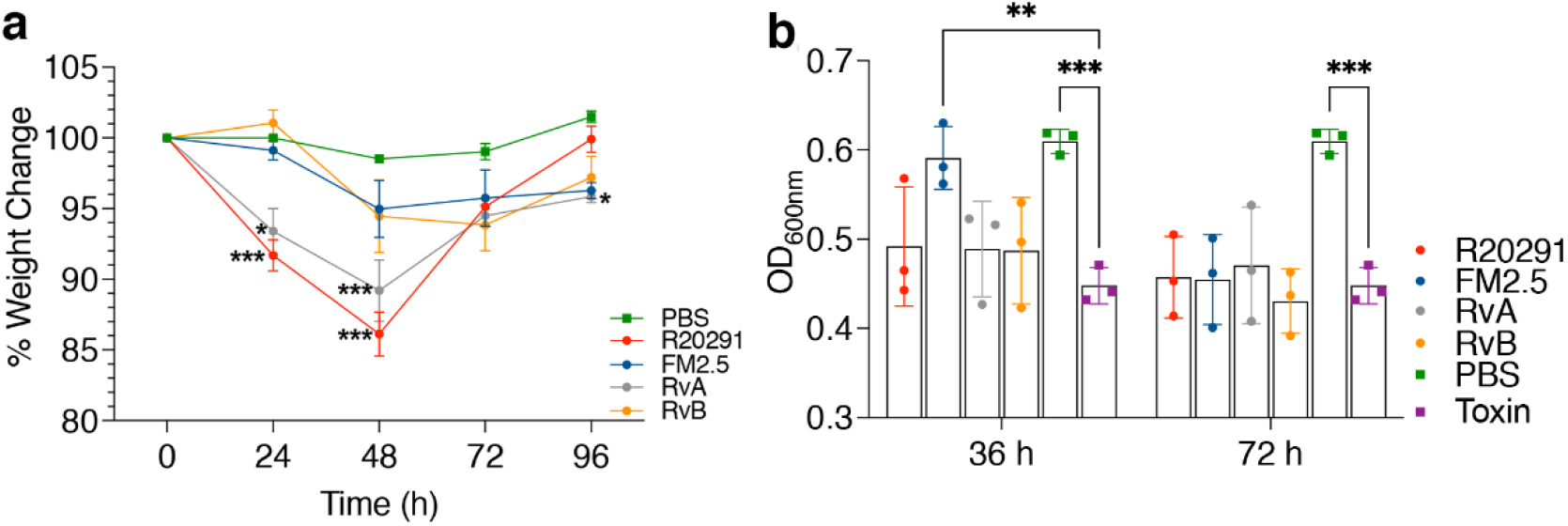
Functional analysis of RvA and RvB *in vivo* and *in vitro*. (**a**) Female C57/Bl6 mice were challenged with spores of R20291, FM2.5, RvA, RvB or mock infected with sterile PBS. Weight loss was monitored at the same timepoint each day for four consecutive days following infection. Each point is the average of multiple individuals, with at least 5 animals. (**b**) *In vitro* toxin activity as measured through challenge of Vero cells. Samples were prepared by filtering supernatant following *C. difficile* growth for 36 or 72 h and activity was measured through challenge of Vero cells. Supernatants were harvested at the same phase of growth for each strain. OD_600_ represents the optical density of Giemsa stain incorporated and released from intact Vero cells, hence high OD represents limited toxicity. Results displayed are the mean ± SEM of at least three independent replicates. Statistical tests were conducted using GraphPad Prism software v.12. Statistical tests include one-way ANOVA with Tukeys post-test; or a student’s t-test with Welch’s correction. Statistical significance is indicated: ns – not significant; *p < 0.05; **p < 0.01; and ***p < 0.001.

To determine whether these differences were associated with changes in toxin production, the revertants were cultured *in vitro* and filtered spent growth medium from 36 and 72h growth was used to determine the level of toxin B activity. Both RvA and RvB produced comparable levels of toxin to R20291 at these time points (Fig. 5b), with toxin-mediated damage to Vero cells, resulting in cell rounding, cellular loss and reduced levels of staining with Giemsa. In agreement with previous reports (Kirk et al. 2017), FM2.5 produced less toxin than R20291 at 36 h, although toxin production levels were comparable in all strains by 72 h.

### Modification of the S-layer results in large changes in gene expression

To gain a greater understanding of the differences in gene expression between R20291, FM2.5 and revertant strains, comparative RNA-Seq analysis was conducted following *in vitro* growth. Analysis revealed differences in gene expression between R20291 and FM2.5, with over 287 differentially expressed genes (DEGs) (Fig. 6a), linked to alterations in metabolism, transport, membrane integrity and sporulation (Fig. 6b). In contrast, less differences were observed when R20291 was compared to RvA (44 DEGs), than RvB (185 DEGs), which showed similar numbers of DEGs to FM2.5. This correlates well with the observed behaviour of these strains within animals (RvA associated with WT-like disease and RvB with FM2.5-like attenuation). Analysis of these data suggest that differences in observed disease severity could be linked to changes in transcription of several virulence-associated traits, including toxin A and B, and genes associated with sporulation. While the recovery of an intact S-layer would appear sufficient in the case of RvA, to restore wild type gene transcription, only partial transcription profile restoration, including toxin expression, is observed in RvB. However, as toxic activity was observed at 36h in culture, the alterations in transcription control seen in FM2.5 and RvB would appear to be limited to timing rather than absolute prevention of toxin production.

**Fig. 6.**
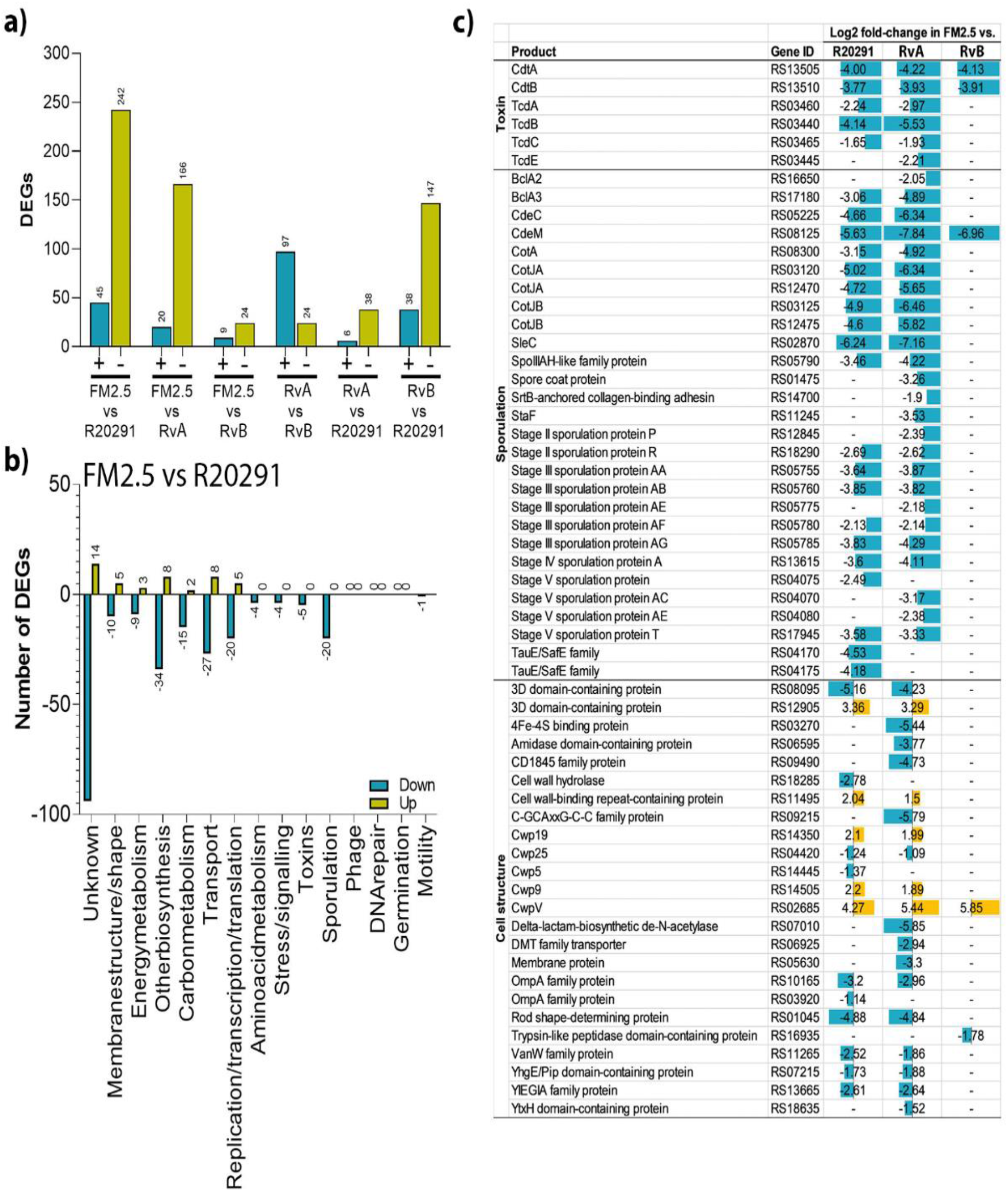
Global transcriptional differences between isolates of *C. difficile* following *in vitro* growth. Analysis of mRNA recovered from *in vitro* grown cultures of *C. difficile* isolates R20291, FM2.5, RvA and RvB. (**a**) Total number of differentially expressed genes (DEGs) between experimental groups are highlighted in blue (upregulated) and gold (downregulated). (**b**) DEGs from experimental comparison of R20291 and FM2.5 were categorised based on function. (**c**) Transcriptional differences in select genes of FM2.5 relative to experimental comparison with R20291, RvA and RvB.

Taken together, these data suggest that expression of the S-layer plays a key role in *C. difficile* disease within the host.

## Discussion

The S-layer of *C. difficile* has long been considered as integral to its physiology and pathogenesis, with several roles reported, including adherence to the epithelial barrier (Merrigan et al., 2013), immune cell signalling (Ryan et al., 2011; Chen et al., 2020), sensitivity to antimicrobial peptides (Kirk et al. 2017) and sporulation efficiency (Kirk et al. 2017). Here, we describe the pathogenesis of the S-layer-null mutant FM2.5 within the mouse model of disease and report the recovery of toxin-independent, spontaneous S-layer variants in which SlpA expression is restored. This unexpected but reproducible phenomenon supports the growing evidence that this structure plays a key role in adaptation and survival within the host.

FM2.5, a strain originally selected through its resistance to the R-type bacteriocin Av-CD291.2, was previously reported to be attenuated in the Syrian hamster model of *C. difficile* (Kirk et al. 2017). These studies indicated that SlpA was essential for disease, with no diarrhoeal symptoms observed in infected animals, despite the recovery of FM2.5 from the caecum and colon of infected hamsters 14 days pi. While here we confirmed that the attenuated phenotype was reproducible in mice, we also detected SlpA revertant clones from infected animals. Interestingly, SlpA revertants were not observed during hamster infections, despite using the same chromogenic agar for recovery of FM2.5 isolates from infected animals.

Isolation of FM2.5 SlpA revertants in mice but not in hamsters suggests that differences in the local environmental conditions between the hamster and mouse may influence the amplification and outgrowth of these S-layer variant strains. Indeed, it has been suggested that *C. difficile* colonisation efficiency may reflect variation in expression of cathelicidins (such as LL-37) within these hosts (Woods et al. 2018), which would help to explain why revertants were not amplified in the hamster gut. This observation also lends support to the hypothesis that an intact S-layer confers resistance to the antimicrobial activity of enzymes (such as lysozyme) and antimicrobial peptides (LL-37) (Kirk et al. 2017). The observation that FM2.5 is acutely sensitive to LL-37 and lysozyme (Kirk et al. 2017) and becomes resistant following S-layer restoration (Lanzoni-Mangutchi, 2022) further supports the premise that revertants with an intact S-layer have a competitive advantage *in vivo*.

Selection and amplification of the revertants in this study further highlights the competitive advantage offered by the S-layer in *C. difficile* intestinal survival. Poor recovery of *C. difficile* at 24 hpi in FM2.5-infected mice could also be linked to lower or less efficient rates of germination by FM2.5 compared to R20291. However, *in vitro* studies using taurocholic acid as a germinant, indicate that these strains show equivalent rates of germination (Kirk et al. 2017). Alternatively, lower rates of FM2.5 at 24h hours could be a result of an increased susceptibility of the SlpA-null mutant to anti-microbial peptides within the gut. The equivalent recovered numbers at 48 and 72h for infections with FM2.5 when compared to R20291 may correspond to an increased population of revertant clones, considering the high level of reversion, as quantified by genetic analysis.

Although SlpA variants have not been observed during sequential growth of the FM2.5 isolate *in vitro*, several factors may influence their presence, albeit at low numbers, within the inoculum used to infect mice. As a largely obligate anaerobe, the sensitivity of *C. difficile* vegetative cells to oxygen complicates their use in animal models of infection. In contrast, the preparation and use of spores correlates with the natural infection and avoids co-administration of toxins expressed during *in vitro* growth. However, as FM2.5 has a known reduced sporulation efficiency (Kirk et al. 2017), it is possible that any variants expressing an intact S-layer present would sporulate more efficiently and therefore represent a higher proportion of the inoculum used for infection. Sequence analysis of *slpA* of several different batches of FM2.5 spores supported this hypothesis, with a small (>2.5%) proportion of the population displaying variations upstream of the original *slpA* mutation.

A delay in toxin production by FM2.5 has been reported previously (Kirk et al. 2017) which, coupled with the apparent delayed growth *in vivo*, could account for the difference in weight loss between R20291 and FM2.5 infected mice after 24 hpi. Using our FM2.5*ΔPaLoc* strain, we were able to demonstrate that the weight loss is entirely dependent on toxin production. However, this raises the question as to why animals infected with FM2.5, that show high and equivalent levels of tissue colonisation by 48 hpi, do not show equivalent levels of weight loss and tissue inflammation as R20291 infected animals. Several studies have shown that the S-layer is essential in immune activation (Jarchum et al. 2012; Mamareli et al. 2019), driving the production of proinflammatory cytokines via TLR4/MyD88 dependent pathways and enhancing the toxin-activated inflammasome (Ryan et al., 2011; Cowardin et al., 2015; McDermott et al., 2016). Together, this implies that the timing or spatial localisation of toxins and the S-layer relative to the epithelial barrier could be crucial to immune activation. This is further supported by the observation that RvB showed an equivalent reduction in toxin expression to FM2.5 in the RNA-Seq analysis and a reduction in disease severity in the mouse.

Alternatively, as a feedback mechanism between sporulation and the complex regulatory network controlling toxin production has been proposed (Deakin et al. 2012), it is possible that a defect in FM2.5 sporulation may result in a delay in toxin production. The observed reduction in toxin activity at 36h in the filtered culture supernatant from FM2.5 correlated well with the log2 fold reduction in *tcdB* transcripts (−4.14) and *tcdA* (−2.24) in mRNA recovered from FM2.5 cultures grown for 6h, when compared to transcripts recovered from R20291 cultures, at the equivalent timepoint. In contrast, RvB which also presented a reduction in toxin gene expression compared to R20291 by RNA-Seq, demonstrated equivalent toxin functional activity to R20291 when grown *in vitro* for 36h; RvA and R20291 showed comparable toxin expression in both systems. This suggests that whilst modifications in S-layer can result in delays to toxin production, these changes do not prevent or limit final activity supporting the hypothesis that disease severity might be linked to the timing and co-ordination of S-layer and toxin by the host.

Mice infected with either RvA or RvB showed different disease severity, as indicated by differences in weight loss. The low virulence of an RvB infection may be, in part, explained by the structural differences. Indeed, structural analysis of SlpA_RvB_ revealed a new packing of SlpA molecules in the array, with a rearrangement of the position of SLP_H_ and both interacting domains, now involved in tiling of the S-layer (Fig. 3). This suggests a considerable degree of adaptability of both SLP_L_, where the reversion is located, and SLP_H_, to accommodate varying interactions between neighbouring molecules. The absence of density to model D2 in SlpA_RvB_ further illustrates that this domain is dispensable for S-layer assembly, as previously reported (Lanzoni-Mangutchi et al., 2022). As S-layer assembly is maintained mostly by hydrogen bonds and salt bridges (Lanzoni-Mangutchi et al., 2022), rearrangement of the subdomains to create structures with complementary surface charges seems to enable the different assemblies observed so far. These changes in quaternary structure, with minor changes of secondary and tertiary structure of the subdomains, suggests that the ability to form a paracrystalline array is central to S-layer function and can be achieved in different arrangements. The S-layer must retain a certain degree of flexibility, not only to account for the cell pole curvature and allow cell division, but also for incorporation of minor cell wall proteins that enhance functionality. The presence of a more intricate network of interactions and more extensive interface areas when compared to the R20291-related SlpA_RΔD2_ structure (Table S2) between neighbouring molecules seen in the SlpA_RvB_ structure suggests a potentially less flexible paracrystalline array, with less ability to incorporate specific functions of minor CWPs, which may help to explain differences in disease patterns observed between RvB and the other revertant strain, RvA; the structure of which needs to be studied in more detail. Further structural studies of SlpA_RvA_ revertant and SlpA_R20291_ as well as detailed analysis of S-layer assembly and composition, including the capacity to incorporate other minor cell wall proteins, will help elucidate the role of specific aspects of the S-layer.

While identification of SlpA revertants was unexpected and adds complexity to the interpretation of the data from the mouse disease model, the rapid recovery of these strains highlights the key contribution that the expression of an intact S-layer offers to *C. difficile* infection *in vivo*. This work supports previous observations that strains lacking the S-layer are less virulent *in vivo*, although it remains difficult to identify the specific contribution of the S-layer in the infection process. Instead, this work highlights the potential multifunctional contribution that the S-layer plays in disease as, despite the number of differentially expressed genes observed between R20291, FM2.5 and the revertants *in vitro*, recovery of the intact S-layer was sufficient to restore virulence, at least in one revertant. Importantly, isolation and characterisation of these variants, together with greater knowledge of gene regulation and metabolic pathways impacted, offers a new opportunity to better understand the role of the S-layer in *C. difficile* pathogenesis.

## Materials and Methods

### Bacterial strains and growth conditions

The bacterial strains used in this study include *C. difficile* strain R20291, its derivative FM2.5 (Kirk et al. 2017), RvA, RvB, RvC, RvD and FM2.5Δ*PaLoc* (this study). Strains were routinely grown under anaerobic conditions on Braziers cycloserine, cefoxitin egg yolk (CCEY) agar (Oxoid, UK); CHROMID® *C. difficile* Chromogenic medium (bioMérieux); or in Tryptone yeast (TY) broth (Oxoid, UK).

### Generation of FM2.5Δ*PaLoc*

Homologous recombination was used to generate a derivative of strain FM2.5 that lacked the entire pathogenicity locus (PaLoc). Briefly, 1.2 kb up and downstream of the PaLoc was amplified by PCR using RF920 (cgtagaaatacggtgttttttgttaccctaTGGAATTTAGATATAAAAACCAATTC) and RF921 (atttattttggtgtgGACAACATTGGAATTAAATCAG), and RF922 (aattccaatgttgtcCACACCAAAATAAATGCC) and RF923 (gggattttggtcatgagattatcaaaaaggCCCAACTATGGAAAAACC), respectively, and cloned by Gibson assembly into plasmid pJAK112 (Fuchs et al. 2021) that had been linearised by PCR using RF311 (TAGGGTAACAAAAAACACCG) and RF312 (CCTTTTTGATAATCTCATGACC). The resulting plasmid, pJAK143, was then conjugated into *C. difficile* (Kirk and Fagan 2016) and mutagenesis to knock out the PaLoc was carried out using standard allele exchange (Cartman et al. 2012b).

### Murine model of infection

All procedures were performed in strict accordance with the Animals (Scientific Procedures) Act 1986 with specific approval granted by the Home Office, UK (PPL 60/8797 and PPL PI440270). Food and water were provided *ad libitum* and animals kept at a constant room temperature of 20–22 °C with a 12 h light/dark cycle. Groups of up to six C57/bl6 mice aged 6–8 weeks supplied by Charles River (Edinburgh) were used in each treatment group. An antibiotic cocktail (kanamycin [0.40 mg ml^−1^]; metronidazole [0.215 mg ml^−1^]; colistin [850 U ml^−1^]; gentamicin [0.035 mg ml^−1^]; and vancomycin [0.045 mg ml^−1^] [all Sigma Aldrich, UK]) was administered *ad libitum* in the drinking water as previously described (Jukes et al., 2020) with clindamycin sulphate (150 mg Kg^−1^), administered by oral gavage following cessation of the antibiotic cocktail. Animals were each challenged with approximately 10^5^ spores of *C. difficile* 72 h after clindamycin treatment. Mice were monitored closely post-infection and weighed daily to determine the severity of the disease. Animals with a weight loss greater than 10 % of pre-challenge weight were given soft food and were culled if weight loss reached 20 %.

### *C. difficile* shedding and organ colonization

Fresh faecal samples collected daily were weighed, serially diluted in phosphate buffered saline (PBS) and cultured on CCEY agar at 37 °C for 48 h. At the experimental endpoint, animals were culled, and the caecum and colon harvested. Enumeration of total counts and spore-specific counts in lumen associated material were performed as previously described (Jukes et al., 2020). In brief, total viable counts were determined by plating serial dilutions on ChromID selective media (Biomeuriex). Spores were enumerated following heat treatment at 56 °C for 20 min.

### Quantification of toxin expression

Quantification of toxin activity was performed using monolayers of Vero cells (kidney epithelial cells) as described previously (Buckley et al. 2011). Briefly, toxin was recovered from the spent filtered TY medium used to support bacterial growth for 36-72 h. Spent medium was recovered at the same stage of the growth cycle and at the same OD_600nm_. *In vivo* toxin activity was measured by filtering luminal content collected from the caecum and colon of infected mice. Luminal content collected from the caecum and colon of uninfected mice was used as a control. Samples for toxin measurement were tested by the addition of serial dilutions to confluent monolayers within 72 h of collection. Cells and toxin were co-cultured for 24 h before cells were washed with phosphate buffered saline (PBS), fixed with 5 % formal saline (Fisher), and stained with Giemsa for 30 min, before thorough washing to remove excessive stain. For data presented in Figures 1d, and S3, toxin activity was determined as the reciprocal of last dilution in which toxin activity was observed, i.e. showing cell destruction. In Figure 5, in an attempt to quantify the toxin activity more precisely, excess stain was removed by washing with PBS before cells were permeabilised to release internalised stain using 200 μl 1% SDS. 100 μl of the supernatant was transferred to a fresh plate and the OD_620nm_ values determined. In this context higher values indicate Vero cells are intact and unaffected by the toxin, lower values indicate that toxin mediated damage prevents uptake and retention of the dye.

### Histology and immunohistochemistry

Tissue samples were harvested from the caecum and colon of antibiotically susceptible animals infected with either R20291 or FM2.5 at post-mortem, 48 hpi. These tissues were gently washed in sterile PBS and immediately fixed in 10% formalin. Embedded tissue sections were cut and stained with Hematoxylin and Eosin (Jukes et al., 2020). Blind histological scoring of tissue was performed on 3 independent sections of caecal and colonic tissue. Each section was scored out of a total of 20, with a score of 1 indicating no change, 2 mild change, 3 moderate change and 4 severe changes, for the following categories: epithelial damage, neutrophil migration, haemorrhagic congestion, tissue oedema and crypt hyperplasia. Data presented represents the mean scores for 3 mice for each treatment.

### *slpA* sequencing from isolated revertant clones

Individual clones of bacteria, recovered from faecal or tissue associated material that showed different morphology on ChromID plates, where subject to at least two rounds of clonal selection. Genomic DNA was isolated from a 20 ml culture grown anaerobically for 18 h in tryptic soya broth (TSB). Bacterial cells were initially disrupted enzymatically by resuspending the pellet in lysis buffer (20 mM Tris-Cl, pH8.0, 2 mM Na EDTA, 1.2 % Triton X-100, lysozyme 200 mg ml^−1^), and incubated at 56 °C for 90 min. The DNA was recovered using the DNeasy Blood and Tissue Kit (Qiagen), following manufacturer’s instructions. A 478 bp fragment of *slpA*, centred on the FM2.5 point mutation (Kirk et al. 2017), was amplified by using oligonucelotides RF110 (GACATAACTGCAGCACTACTTG) and RF111 (CAGGATTAACAGTATTAGCTTCTGC). The resulting fragments were subjected to Sanger sequencing and compared to wild type and FM2.5 sequences.

### Isolation and sequencing of *slpA* from faecal extracts

Faecal samples were also collected for sequencing of *slp*A, by directly extracting DNA from faecal samples using the FastDNA SPIN kit for soil (MP Biomedicals). Briefly, approximately 200-600 mg of faeces sample was suspended in 978 μl sodium phosphate buffer with 122 μl MT™ buffer lysis solution. Samples were then homogenised in a FastPrep instrument using two 30 second pulses, at speed setting 6.5, in lysing matrix E. Samples were centrifuged for 10 min at 14,000 *x g* to remove debris. 250 μl protein precipitation solution was added to the lysate supernatant, and the precipitant formed removed by centrifugation at 14,000 *x g* for 5 min. DNA was then bound to a silica matrix, washed using the kit wash buffer, and eluted with water.

DNA extracted from faeces was used as a template for PCR amplification of a 330 bp fragment of *slpA* using Phusion polymerase (NEB) and RF2193 (ACACTCTTTCCCTACACGACGCTCTTCCGATCTCTACTTGTAGCTACTTTTA TTGCAC) and RF2194 (GACTGGAGTTCAGACGTGTGCTCTTCCGATCT CAAGGATATACAGTAGTACAGAGC) oligonucleotides. Resulting DNA fragments were purified and sequenced using the amplicon-EZ service offered by GENEWIZ (Azenta Life Sciences).

### Extraction and western immunoblot analysis of S-layer and associated proteins

Surface layer proteins were extracted using low pH glycine as previously described (Fagan et al. 2009) and analysed by SDS-PAGE using standard methods (Laemmli 1970). Proteins were transferred to nitrocellulose membranes via semi-dry transfer (Bio-Rad Trans Blot Turbo; 25 V, 30 min) for western immunoblot analysis. Transfer efficiency was confirmed by PonceauS staining of membrane post transfer, and Coomassie staining of the polyacrylamide gel following transfer (Fig. S5). Membranes were blocked for 1 h in Phosphate-buffered Saline containing 0.1 % Tween20 (PBS-T) with 5 % milk powder. Blots were subsequently incubated in primary antibody (rabbit anti-SLP_H_ raised against *C. difficile* 630 1:100,000 dilution; rabbit anti-SLP_L_ raised against *C. difficile* R20291 1:200,000 dilution) in PBS-T containing 3 % milk powder, for 1 h at room temperature. Membranes were washed thoroughly in PBS-T before incubation with secondary antibodies (anti-rabbit horseradish peroxidase, Promega WB401B 1:2,500 dilution) for 1 h at room temperature. Blots were washed in PBS-T before detection by chemiluminescence (Bio-Rad). Molecular weight (MW) markers (Thermo Scientific™ 26616) were imaged (Bio-Rad ChemiDoc XRS+) simultaneously and overlaid onto the blots to aid visualisation.

### Protein purification and X-ray crystallography

*C. difficile* revertant strains were cultured in 400 ml of TYG broth for 16 h. Cultures were then centrifuged at room temperature at 4,696 x *g* and resulting pellets were washed with 40 ml of 0.01 M HEPES pH 7.4 and 0.15 M sodium chloride (HBS) buffer. S-layer extraction was performed by resuspending the washed pellet in 4 ml of 0.2 M glycine-HCl pH 2.2 and centrifugation for 5 min at 21,100 x *g*. Collected supernatant was then neutralized with 2 M Tris-base. S-layer extract was filtered and resolved onto a Superdex 200 26/600 column using an ÄKTA Pure FPLC system (Cytiva) in 50 mM Tris-HCl pH 7.5, 150 mM NaCl buffer.

Purified SlpA_RvB_ at 10 mg ml^−1^ was subjected to crystallization using a Mosquito liquid handling robot (TTP Labtech), with the sitting drop vapor-diffusion method, at 20 °C. Crystals were obtained in 0.03 M magnesium chloride hexahydrate; 0.03 M calcium chloride dihydrate, 0.12 M ethyleneglycol, 0.05 M Tris (base); 0.05 M bicine pH 8.5, 20% v/v glycerol; 10% w/v PEG 4,000.

Data were collected on the I24 (λ = 0.71 Å) beamline at the Diamond Light Source Synchrotron (Didcot, UK; mx24948-136) at 100 K. The data were acquired from the automatic multi-crystal data-analysis software pipeline xia2.multiplex (Gildea et al. 2022) within the Information System for Protein Crystallography Beamline (ISPyB), re-processed using Automatic Image Processing with Xia-2 (DIALS [Winter et al. 2018] and Aimless 3d [Evans and Murshudov 2013]) and scaled with Aimless within ccp4.cloud of CCP4 (Winn et al. 2011) software suit.

The initial model of the core SLP_H_ was obtained by molecular replacement in Phaser (McCoy et al. 2007), using an SlpA_RvB_ model of CWB2 domains, derived from the SlpA_RΔD2_ model (PDB ID: 7ACZ) and calculated using SWISS-MODEL (Waterhouse et al. 2018). The generated solution model was then subjected to automatic model building with Modelcraft (Cowtan et al. 2020), followed by manual building with Coot (Emsley and Cowtan 2004) and refinement in Refmac5 (Murshudov et al. 2011).

Final models were obtained after iterative cycles of manual model building with Coot and refinement in phenix_refine (Liebschner et al. 2019). Data collection and refinement statistics are summarized in Table S1.

PDBePISA (Krissinel and Henrick 2007) was used to investigate interdomain and protein-protein interfaces in the crystallographic lattice to identify interacting residues, which were confirmed by manual inspection within COOT.

Structural representations were generated using PyMOL Molecular Graphics System (Schrödinger, LLC).

### Protein structure prediction

Homology models for SlpA_R20291_ and SlpA_RvA_ were generated by providing SWISS-MODEL webserver with the SlpA_RvB_ (PDB ID: 8BBY) or SlpA_RΔD2_ (PDB ID: 7ACZ) as user templates, as well as without a template. Structural alignments between predicted models and templates were performed using COOT (Emsley and Cowtan, 2004). As SlpA_RΔD2_ lacks the D2 domain, predicted models based on this experimental model have a disordered D2 domain. Therefore, overall comparison of the three predicted models was based on models calculated in the default mode, which uses SlpA_R7404_ (PDB ID: 7ACX) as a template, while analysis of the reversion-containing region in the D1 domain was done using SlpA_RΔD2_ or SlpA_RvB_ derived models.

### Recovery of mRNA for RNA sequence analysis

RNA was recovered from *C. difficile* strains R20291, FM2.5, RvA and RvB which had been cultured *in vitro* in TY broth. Briefly, bacterial cells reaching an OD_620nm_ = 0.6 were pelleted (5000 x *g*, 15 min) and immediately fixed in 1.5 ml RNA-protect (Qiagen) for 10 min before being processed using a PureLink RNA mini kit (Ambion) to extract total RNA. To ensure maximal lysis of bacteria and recovery of RNA, the bacterial pellet was additionally subject to treatment with 100 ml lysozyme solution (10 mg ml^−1^ in 10 mM Tris-HCl [pH8.0] 0.1 mM EDTA), 0.5 ml 10% SDS solution and 350 ml of Lysis Buffer (Invitrogen PureLink RNA Mini Kit) containing 2-mercaptoethanol. Cells were then homogenised using MP Biomedical beads (0.1 mm) and bead beater (MP Biomedical FastPrep24) with cells subject to 2 cycles of 60 s beating, followed by incubation on ice for 2 min. Total RNA was then extracted using the standard PureLink RNA mini kit protocol, according to the manufacturer’s specifications. Genomic DNA was removed using a TURBO DNase kit (Ambion) and samples were tested for efficient removal of DNA by conventional PCR. Samples for RNA-Seq were prepared in triplicate on two separate occasions (6 samples for each) for all four bacterial strains.

Illumina library preparation of mRNA samples for RNA-Seq was prepared using a TruSeq Stranded mRNA library prep kit (Illumina) according to the manufacturer’s instructions. Sequencing was performed on the Illumina NextSeq 500 platform (75 bp length; single-end). Library generation, optimisation of amplification and sequencing were performed at the University of Glasgow Polyomics facility. Quality control of sequencing data was performed using FastQC (Babraham Bioinformatics) to assess the minimum Phred threshold of 20 and potential data contamination. The raw data will be deposited to the Gene Expression Omnibus reference ID GSE205747.

### RNA analysis and identification of differentially expressed genes

Raw RNA-Seq datasets were subject to the following pipeline. Firstly, fastQ files were assessed using FastP (Chen et al. 2018) and then were aligned to the *C. difficile* R20291 (accession number NC_013316) reference genome using STAR (Dobin et al. 2013) (v2.6) with –quantMode GeneCounts –outFilterMultimapNmax 1 and –outFilterMatchNmin 35. We used a Star index with a –sjdbOverhang of the maximum read length – 1. Next, read count files were merged and genes with mean of < 1 read per sample were excluded. Finally, the expression and differential expression values were generated using DESeq2 (Love et al. 2014) (v1.24). For differential comparisons, we used an A versus B model with no additional covariates. All other parameters were left to default.

The processed data was then visualised using Searchlight (Cole et al. 2021), specifying one differential expression workflow for each comparison, an absolute log2-fold cut-off of 1 and adjusted *p* of 0.05. All other parameters were left to default.

### Pathway Analysis Methods

Functional and metabolic pathways were implied by interrogation of the WP numbers assigned using the *C. difficile* annotated genome (NC_013316.1) and entered into Uniprot or NCBI Blastp databases. Gene ontology (GO) was assigned based on Biological Process assignment within Uniprot.

### Statistical analysis

Statistical analysis was carried out in GraphPad Prism v.9. The tests and parameters used are detailed in the figure legends throughout. Tests used included *t*-test with Welch correction, ANOVA with Tukey’s post-test and Kruskal–Wallis with Dunn’s multiple comparisons.

## Acknowledgments

This work was supported by the Wellcome Trust [204877/Z/16/Z]. FV, MO, ABS, JAK and JH were supported by this Wellcome Trust Collaborative Award, awarded to GRD, PSS, RPF. The authors would like to thank Dr Arnaud Baslé, facility manager of the Newcastle Structural Biology Lab, and beamline staff at I24, Diamond Light Source (BAG mx24948) for support on structural data collection. The contents of this work are solely the responsibilities of the authors and do not reflect the official views of any of the funders, who had no role in study design, data collection, analysis, decision to publish, or preparation of the manuscript.

## Contributions

MO, FV, GRD carried out animal experiments, collected and analyzed data, wrote and revised the manuscript. JAK designed and performed experiments to generate the FM2.5Δ*PaLoc* strain. JAK, RRC and RFP carried out analysis on the sequential samples from the mouse experiments, wrote and revised the manuscript. ABS, PLM and PSS carried out crystallization, structural determination, modelling and analysis of the revertants, wrote and revised the manuscript. MO, JH and JC carried out the transcriptomics analysis, MO and JH processed the samples and JC undertook bioinformatic analysis. GRD, PSS and RPF designed experiments, analyzed the data, supervised the study, wrote and revised the manuscript.

## Conflict of Interest

The authors have no competing interests that might be perceived to influence the interpretation of the article.

## Notes

### Competing Interest Statement

The authors have declared no competing interest.

